# Cannabidiol Toxicity Driven by Hydroxyquinone Formation

**DOI:** 10.1101/2024.10.22.619647

**Authors:** Metzli I. Montero, Pravien S. Rajaram, Jose E. Zamora Alvarado, Kara E. McCloskey, Ryan D. Baxter, Roberto C. Andresen Eguiluz

## Abstract

Oxidative byproducts of cannabidiol (CBD) are known to be cytotoxic. However, CBD susceptibility to oxidation and resulting toxicity dissolved in two common solvents, ethanol (EtOH) and dimethyl sulfoxide (DMSO), is seldom discussed. Furthermore, CBD products contain a wide range of concentrations, making it challenging to link general health risks associated with CBD cytotoxicity. Here, we report on the effect of CBD and CBD analogs dissolved in EtOH or DMSO at various concentrations. The cells used in these studies were human umbilical vascular endothelial cells (HUVECs). Our findings show that significant CBD oxidation of CBD to form cannabidiol-quinone (CBD-Q) and subsequent cytotoxicity, occurring at 10 µM concentration regardless of the solution delivery vehicle. Moreover, a new analog of CBD, cannabidiol-diacetate (CBD-DA), exhibits significantly more stability and reduced toxicity compared with CBD or CBD-Q. This knowledge is important for determining concentration-dependent health risks of complex cannabinoid mixtures and establishing legal limits.

## INTRODUCTION

Cannabis has been the source of social and political debate for decades, perpetuated by its contradictory therapeutic and detrimental effects on human health. A few therapeutic uses include the treatment of mental health disorders, chronic pain management, cancer treatment, and alleviating chemotherapy-induced nausea.^1–5^ However, detrimental effects include increased susceptibility to respiratory diseases and adverse cardiovascular events, such as bronchitis and stroke.^6–10^ As a result of conflicting results, worldwide policies regulating cannabis use are highly varied. Research aiming to understand the efficacy and safety of the over 550 chemicals that have been identified in the plant is growing.^11,12^

One of the chemicals isolated from cannabis is cannabidiol (CBD), a non-psychoactive phytocannabinoid with ongoing investigations in various pharmacological contexts, and one FDA-approved drug, EPIDIOLEX^®^, for treating two epilepsy disorders already in US markets.^13,14^ Current studies are also examining CBD’s pharmacological potential to treat pain and cancer, including its ability to inhibit angiogenesis, attenuate the inflammatory response, and regulate vasodilation and vasoconstriction.^15–24^ To investigate CBD use, researchers have utilized both *in vivo* animal and *in vitro* human cell culture models. However, drug dosage, route of administration, and individual clinical history within specific contexts all play critical roles in the efficacy or harm after administration or consumption, complicating quantitative outcomes assessments. Additionally, there is more variety in drug source, drug vehicle, and sample preparation between current studies, making it even more difficult to compare studies.^25^

To address these ongoing challenges, we present evidence supporting the hypothesis that the toxicity of oxidized cannabinoids contributes to the adverse health effects associated with cannabis use. To test this hypothesis, we first demonstrate that CBD oxidizes to form cannabidiol-quinone (CBD-Q) in a dose-dependent manner in two frequently used solvents in cell culture, ethanol (EtOH) and dimethyl sulfoxide (DMSO). Then, we used these two solvents as drug vehicles for CBD, CBD-Q, and a more stable cannabidiol, cannabidiol-diacetate (CBD-DA), in cytotoxicity studies involving human umbilical vein endothelial cells (HUVECs) to mimic the first cells exposed to CBD of intravenous drug delivery. We compared the effects of both drug vehicle and drug dosage on cell viability. Following a protocol from pre-existing literature, we tested two dosages: 1 and 10 µM.^26,27^

With this study, we confirmed that CBD-Q was more toxic than CBD and CBD-DA, with all analogs presenting concentration-dependent toxicity. Furthermore, our findings support other reports showing that above a critical concentration (as is the case for 10 µM)^26^ leads to the induction of cellular apoptosis. In contrast, we also see proliferative effects at lower concentrations (1 µM), suggesting cell protectivity.^26^ With this investigation, we emphasize CBD’s instability, how this instability may affect toxicity studies, the importance of detailing drug vehicle storage and preparation, and the need for continuing comparative studies involving the impact of drug vehicles on CBD and its analogs.

## RESULTS & DISCUSSION

The stability of CBD was quantified and compared with two additional CBD analogs: an isolated cannabidiol quinone denoted as CBD-Q and a synthesized cannabidiol-diacetate (CBD-DA) (SI S1 Synthesis Procedures). To prevent sample degradation, CBD and its analogs were stored under an 99% argon atmosphere at -20 °C for up to one month prior to cell culture experiments. Upon retrieval, they were dissolved in DMSO or ETOH and used immediately.^28^ For investigating the long-term stability of CBD in solution, CBD in DMSO and CBD in EtOH solutions were also stored in the dark for one month at 4 ºC, and then characterized using mass spectroscopy (SI S2. Mass Spectrometry). After one month, mass spectroscopy revealed that both the DMSO and EtOH samples displayed a decrease in the relative abundance of CBD. While the relative abundance of CBD in EtOH decreased with 20% remaining and 80% converted to CBD-Q, CBD in DMSO solution completely degraded with no trace of CBD after one month with 100% converted to CBD-Q (Figure 1a).

**Figure 1.**
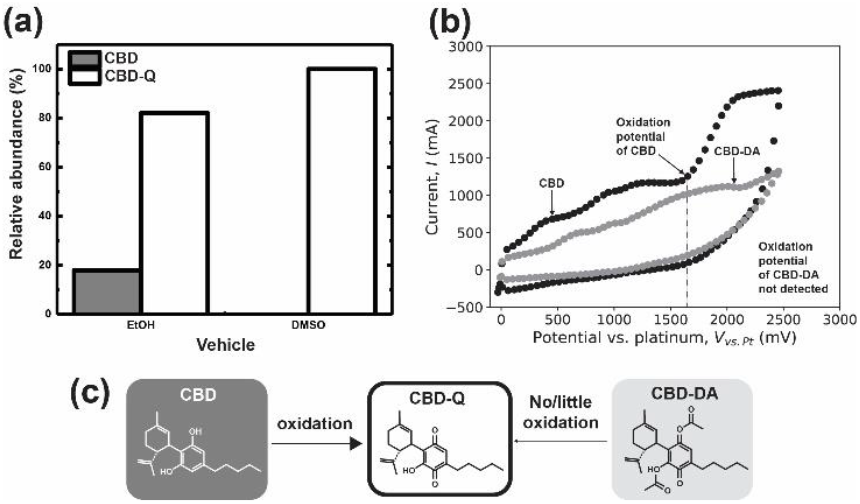
Stability of CBD, CBD-Q, and CBD-DA. (a) Relative abundance of CBD and CBD-Q in dissolved and stored in DMSO or EtOH in the dark for one month, (b) cyclic voltammogram displaying the electron flow for CBD and CBD-DA with oxidation potential of CBD, and (c) CBD oxidation to CBD-Q and lack of oxidation of CBD-DA to CBD-Q.

The 100% loss of CBD in DMSO, and the relative abundance of CBD-Q in both samples indicate that oxidation of CBD to CBD-Q is greater in DMSO. Cyclic voltammetry was performed to assess the oxidative susceptibility of CBD and CBD-DA (SI S3. Cyclic voltammetry procedures). The cyclic voltammogram (Fig. 1b) shows a distinct oxidation potential for CBD at approximately 1600 mV, indicating its high oxidative susceptibility. This supports the conclusion that storing CBD in an oxygen-rich environment leads to oxidation products like CBD-Q. In contrast, no significant oxidation potential was observed for CBD-DA analog, suggesting that the hydroxyl groups on CBD are likely involved in the formation of CBD-Q (Fig. 1c). Together with prior research demonstrating that DMSO exhibits a lower oxygen solubility compared with EtOH, factors such as storage temperature and light may also be contributing to the production of oxidation products.^29^ Knowing how CBD may degrade into CBD-Q in solution over time, we used any dissolved CBD analog samples for cell culture studies immediately upon retrieval form storage in -20 °C argon.

For cell toxicity assays, we followed previously described protocols.^26,27^ We diluted the analog-vehicle solutions into EGM-2 (Lonza, CC-3162) cell culture medium and treated HUVECs to working concentrations of 1 and 10 µM for 24 hrs. Afterward, cells were stained with Calcein AM (Invitrogen, C3099) and imaged at 10x using a fluorescent microscope. Images were then analyzed using FIJI with pre-determined size-exclusion thresholds used for cell counting, followed by a student *t*-test to determine statistically significant differences between conditions.^30^ Detailed methods can be found in SI S4. Cell cutlure and S5. Quantitative analysis of cytotoxicity and S6. Statistical analysis.

At the lowest 1 µM concentration of CBD, CBD-Q, and CBD-DA, all cannabinoid analogs yielded a slight decrease in the average live cell count compared to the solution control (Fig 2a and 2b). Still, this decrease was only statistically significant in the EtOH control compared with the 1 µM CBD-DA (Fig 2b).

**Figure 2.**
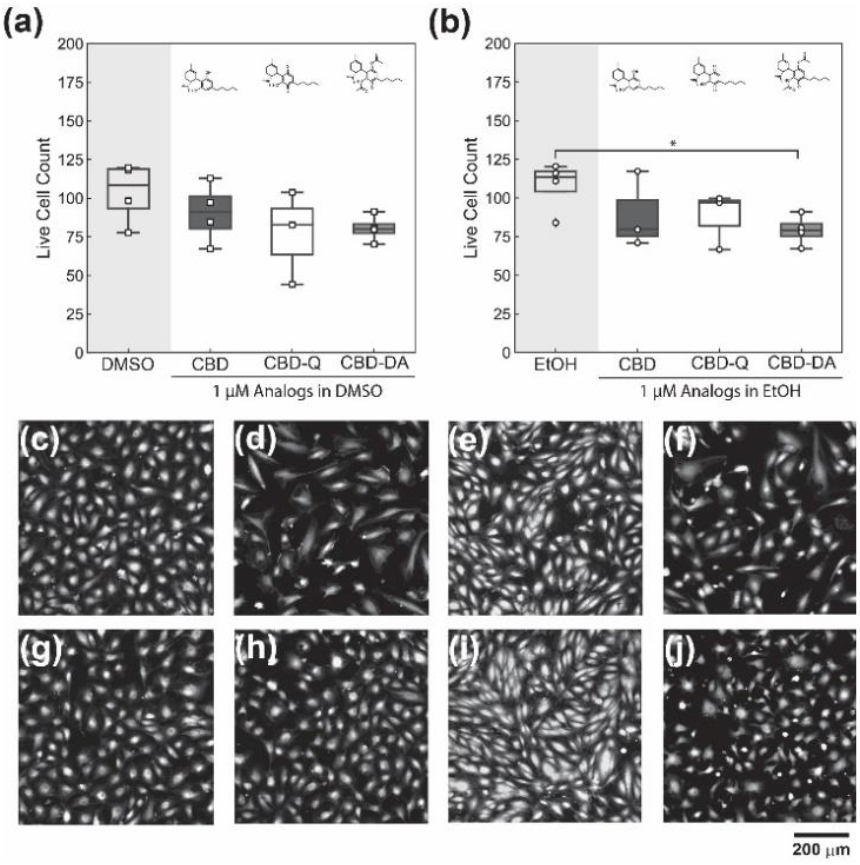
Stability of CBD, CBD-Q, and CBD-DA at 1 µM in (a) DMSO as a vehicle and (b) EtOH as a vehicle. Boxplots include the averaged cell counts of each external replicate, represented by the data points, and whiskers representing the upper and lower quartile. Micrographs of HUVECs with live stain calcein AM exposed to (c) only DMSO (N=4), (d) CBD in DMSO (N=4), (e) CBD-Q in DMSO (N=3), (f) CBD-DA in DMSO (N=3), (g) only EtOH (N=4), (h) CBD in EtOH (N=4), (i) CBD-Q in EtOH (N=3), and (j) CBD DA in EtOH (N=3). Scale bar = 200 µm.

Additionally, the endothelial cell morphologies confluency appears slightly different in several conditions compared to the controls, but cell morphology differences are likely related to confluency in each image.

Because the toxicity of the oxidized metabolites may not directly correlate to the oxidation potentials, it is essential to establish which cannabinoids yield products posing the most significant risk for adverse health effects.

At the highest concentration, 10 µM, as hypothesized, all conditions displayed a more significant decrease in the average live cell count than for 1 µM conditions, with the CBD and CBD-Q in DMSO and EtOH exhibiting the largest cytotoxicities. However, for the CBD, CBD-Q, and CBD-DA in DMSO, a student *t*-test analysis indicated all of the 10 µM DMSO conditions possessed an averaged live cell count that was significantly lower than the control (Fig 3a), with 10 µM CBD-DA exhibiting the highest cell survivability, with cell counts at 75 ± 2 per field of view. In contrast, CBD and CBD-Q measured 60 ± 10 and 30 ± 9 cell counts per field of view, respectively. Moreover, 10 µM CBD, CBD-Q, and CBD-DA treatment groups were not statistically different, demonstrating that at 10 µM, all CBD analog conditions displayed significant cytotoxicity.

**Figure 3.**
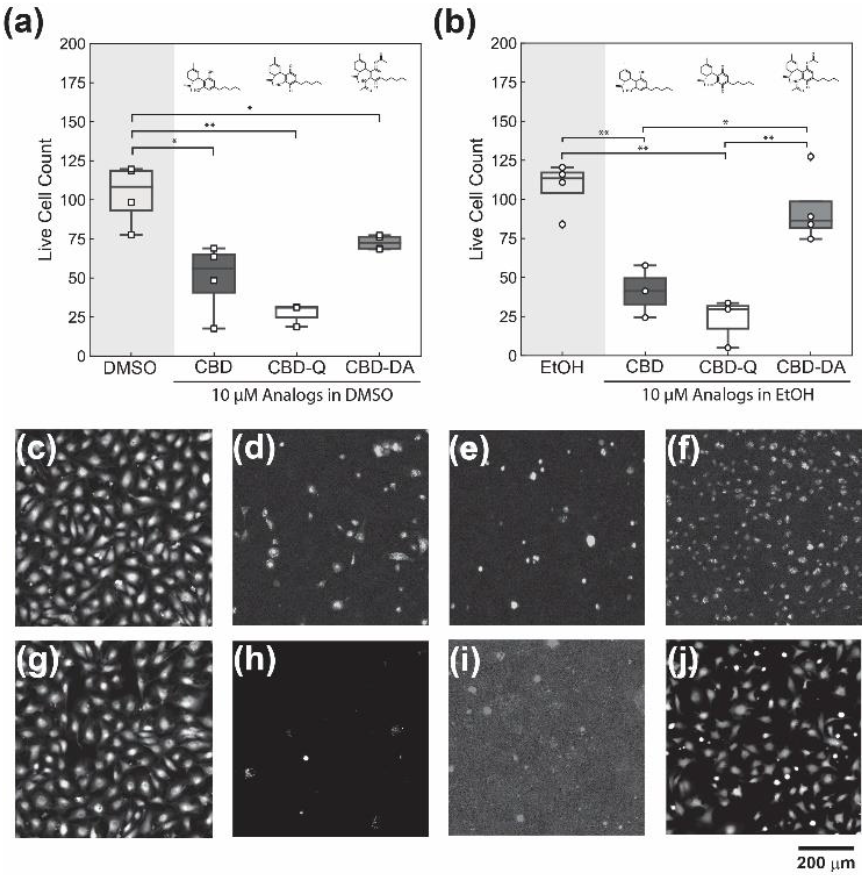
Stability of CBD, CBD-Q, and CBD-DA at 10 µM in (a) DMSO as a vehicle and (b) EtOH as a vehicle. Micrographs of HUVECs with live stain calcein AM exposed to (c) only DMSO, (d) CBD in DMSO, (e) CBD-Q in DMSO, (f) CBD-DA in DMSO, (g) only EtOH, (h) CBD in EtOH, (i) CBD-Q in EtOH, and (j) CBD DA in EtOH. Scale bar = 200 µm.

The analogs delivered in the EtOH vehicle exhibited similar trends, with CBD and CBD-Q in EtOH treatments significantly decreasing cell survivability relative to the EtOH control. However, the average live cell count of CBD-DA in EtOH was only slightly lower than the EtOH control and this decrease was not statistically significant. When comparing the CBD, DBD-Q, and CBD-DA only (not against the control), the cells treated with CBD-DA in EtOH possessed a significantly greater average live cell count per field of view and considerably higher survivability than CBD and CBD-Q in EtOH. The decrease in cell viability in CBD and CBD-Q in both DMSO and EtOH is easily observed visually (Fig. 3 c-j), where a complete eradication of the cell monolayer and overall reduction of cell attachment is evident while the CBD-DA micrographs from both DMSO and EtOH solution treatments (Fig. 3f and Fig. 3j) show that significantly more live cells remain. Still, some differences in cell morphology can be seen when comparing control samples and CBD-DA micrographs. The calcein stain appeared more continuous and localized around the center of the cell body in the controls while in the CBD-DA micrographs, one could observe a slight speckling of the fluorescent signal, suggesting a reduction of fluorescent calcein production, which could result from cell apoptosis. Regardless, the presence of the cell within the micrograph indicates that the cells remained attached after washing with PBS and before imaging after 24 hours. We also observed some edge-effect-dependent toxicity (not shown), where cells at the well edges experienced a greater cell death compared with those at the center of the well. This greater cell death might have been due to the lower cell density at the well-edges, suggesting toxicity also to be cell density-dependent. To mitigate this, all images presented and analyzed were of images taken at the approximate center of the well.

Taking these considerations into account, our results demonstrate that CBD in DMSO or EtOH is unstable and has the potential to degrade into CBD-Q, a significantly more cytotoxic analog oxidatively. Even though EtOH exhibits a higher oxygen solubility, cytotoxic cell assays reveal similar results in the effects of CBD analogs, regardless of vehicle, on HUVEC monolayers.

Based on our observations from cyclic voltammetry and cytotoxic assays, we conclude that CBD-Q is a potential culprit for decreased cell survivability. Within any given cell assay, solvents are exposed to some degree of light, oxygen, and/or heat either during the preparation, storage, or experimental process. Since CBD is sensitive to oxygen, degradability can cause experimental results to be unclear as to whether CBD is toxic or whether byproducts, like CBD-Q, are the true source of toxicity. Since most cytotoxic assays are done within a period of 24–72 hours in an open-air environment, it is recommended to reduce any sample exposure to an oxygen-rich environment and preserve sample purity for future cell experiments.

We recommend a few sample storage methods to minimize the oxidation of solvents. In our case, CBD was kept as a crystalline powder, where the powder was pumped and purged to remove air and moisture and then kept in a dark, inert argon environment. Additionally, we used prepared analog solvents immediately, as preliminary results had revealed an increased cytotoxicity depending on the length of analog-solvent storage. This suggests that future cytotoxicity studies on CBD must distinguish whether CBD’s toxicity arises from byproducts produced by CBD or due to the compound’s inherent toxicity. The sensitivity of our samples also highlights the importance of detailing the exact storage conditions and durations of their analog-solvent solutions.

We conclude by emphasizing how new CBD analogs, such as the newly synthesized CBD-DA, can be designed with improved oxidation resistance and reduced cytotoxicity

## CONCLUSION

Understanding CBD analog stability in various environments is vital for cell applications and further studies in drug delivery. Here, we demonstrate that CBD degrades into a cytotoxic compound, CBD-Q, that is increasingly toxic to cells when exposed to oxygen-rich environments. Therefore, we demonstrate the importance of limiting a sample’s oxygen exposure, detailing sample preparation and storage within the literature, and verifying sample purity before usage to compare current and future toxicity studies on CBD and related analogs.

## ASSOCIATED CONTENT

### Supporting Information

Details on experimental approaches and additional schematics and data are provided as a PDF containing:

**S1**. Synthesis Procedures.

**Figure S1**. ^1^H NMR of cannabidiol quinone.

**Figure S2**. ^13^C NMR of cannabidiol quinone.

**Figure S3**: ^1^H NMR of Diacetyl Cannabidiol.

**Figure S4**: ^13^C NMR of Diacetyl Cannabidiol.

**S2**. Mass spectrometry.

**Figure S5:** Mass spectrogram of XYZ.

**Figure S6:** Mass spectrogram of XYZ.

**S3**. Cyclic voltammetry procedures.

**Figure S7:** Cyclic voltammograms of CBD in rich and poor oxygen environments.

**S4**. Cell Culture

**S5**. Quantitative analysis of cytotoxicity

**S6**. Statistical analysis

## AUTHOR INFORMATION

### Corresponding Authors

Ryan Baxter, Department of Chemistry and Biochemistry, University of California, Merced, California 95344, United States;

Roberto Andresen Eguiluz, Department of Chemical and Materials Engineering, University of California, Merced, California 95344, United States; Health Sciences Research Institute, University of California, Merced, California 95344, United States;

## Author Contributions

The manuscript was written with contributions from all authors. All authors have approved the final version of the manuscript. ‡These authors contributed equally.

## ACKNOWLEDGMENT

M.I.M., R.B., K.E.M., J.E.Z.A., and R.C.A.E. acknowledge funding from the National Science Foundation (NSF)-CREST: Center for Cellular and Biomolecular Machines through the support of the NSF Grant No. NSF-HRD-1547848. R.C.A.E. acknowledges funding from the Tobacco-Related Disease Research Program through the support of the University of California Office of the President Grant No. T31KT1583 awarded to R.C.A.E. Additionally, M.I.M acknowledges funding from NIH G-RiSE I-BioSTeP grant No. T32 GM141862. K.E.M. acknowledges funding from the NSF STC: Cellular Engineering in Mechanobiology (CEMB) Grant No. #1548571.

## ABBREVIATIONS

CBD: cannabidiol
CBD-Q: cannabidiol quinone
CBD-DA: cannabidiol diacetate
EtOH: ethanol
DMSO: dimethyl sulfoxide
PBS: phosphate buffer saline

