## Supporting Information for "Cannabidiol Toxicity Driven by Hydroxyquinone Formation"

**Figure S1.** $^{1}$H NMR of cannabidiol quinone.

**Figure S2.** $^{13}$C NMR of cannabidiol quinone.

**Figure S3**: $^{1}$H NMR of Diacetyl Cannabidiol.

**Figure S4**: $^{13}$C NMR of Diacetyl Cannabidiol.

**S2.** Mass spectrometry.

**Figure S5:** Mass spectrogram of XYZ.

**Figure S6:** Mass spectrogram of XYZ.

**S3.** Cyclic voltammetry procedures.

**Figure S7:** Cyclic voltammograms of CBD in rich and poor oxygen environments.

**S4.** Cell Culture

**S5.** Quantitative analysis of cytotoxicity

**S6.** Statistical analysis

### S1. Synthesis Procedures

1.1 Synthesis of desired (1'R,6'R)-6-hydroxy-3'-methyl-4-pentyl-6'-(prop-1-en-2-yl)-[1,1'-bi(cyclohexane)]-2',3,6-triene-2,5-dione (***Cannabidiol Quinone***)

2-[(1R,6R)-6-Isopropenyl-3-methylcyclohex-2-en-1-yl]-5-pentylbenzene-1,3-diol (CBD) (200 mg, 0.63 mmol), was dissolved in 2.5 mL of Acetone. Then, potassium carbonate (K2CO3) (219.4 mg, 1.59 mmol) was added to this solution. After that, the reaction was stirred for 24 hrs at room temperature. After 24 hrs the reaction mixture was deep yellow color. TLC confirmed the completion of the reaction, and the yellow crude was subsequently purified by column chromatography with hexane/ethyl acetate (v/v = 2/1), to afford (1'R,6'R)-6-hydroxy-3'-methyl-4-pentyl-6'-(prop-1-en-2-yl)-[1,1'-bi(cyclohexane)]-2',3,6-triene-2,5-dione (23 mg, 11 % yield). $^{1}$H NMR (500 MHz, cdcl$_3$) δ 6.99 (s, 1H), 6.44 – 6.34 (m, 1H), 5.14 (s, 1H), 4.58 – 4.50 (m, 2H), 3.79 – 3.67 (m, 1H), 2.76 (ddd, *J* = 13.4, 10.8, 2.9 Hz, 1H), 2.40 (tt, *J* = 7.3, 1.7 Hz, 2H), 2.22 (ddd, *J* = 18.0, 9.0, 3.4 Hz, 1H), 2.00 (dd, *J* = 17.4, 5.6 Hz, 1H), 1.77 (ddt, *J* = 12.9, 5.2, 2.3 Hz, 1H), 1.68 (d, *J* = 2.6 Hz, 3H), 1.60 (s, 1H), 1.54 – 1.48 (m, 2H), 1.36 – 1.31 (m, 4H), 1.25 (s, 3H), 0.92 – 0.88 (m, 3H). $^{13}$C NMR (126 MHz, cdcl$_3$) δ 187.16, 184.05, 151.27, 148.40, 144.52, 134.67, 134.00, 122.75, 122.35, 110.65, 44.67, 35.74, 31.36, 30.45, 29.69, 28.82, 28.13, 27.12, 23.42, 22.34, 18.68, 13.90. IR (FTIR): 829, 1052, 1173, 1377, 1584, 1704, 2856, 2927, 3444 cm-1. MS (ESI): m/z calcd. for $C_{21}H_{28}O_3$ $(M+H)^+$ 329.2111, found 329.2111.

1.2 Synthesis of desired (1'R,2'R)-5'-methyl-4-pentyl-2'-(prop-1-en-2-yl)-1',2',3',4'-tetrahydro-[1,1'-biphenyl]-2,6-diyl diacetate (***Diacetyl Cannabidiol***)

2-[(1R,6R)-6-Isopropenyl-3-methylcyclohex-2-en-1-yl]-5-pentylbenzene-1,3-diol (**CBD**) (942 mg, 3 mmol), was dissolved in 5.5 mL of pyridine. Then, to this solution, acetic anhydride (5.4 g, 52 mmol) was added. After that, the reaction was stirred for 24 hrs at room temperature. After 24 hrs the reaction mixture was confirmed by TLC for completion, and the crude was subsequently purified by column chromatography with hexane/ethyl acetate (v/v = 10/1), to afford (1'R,2'R)-5'-methyl-4-pentyl-2'-(prop-1-en-2-yl)-1',2',3',4'-tetrahydro-[1,1'-biphenyl]-2,6-diyl diacetate (1.18 g, 99 % yield). $^{1}$H NMR (500 MHz, cdcl$_3$) δ 6.72 (s, 2H), 5.21 (d, *J* = 2.7 Hz, 1H), 4.56 (t, *J* = 1.8 Hz, 1H), 4.47 (d, *J* = 2.2 Hz, 1H), 3.52 (ddt, *J* = 10.9, 4.5, 2.2 Hz, 1H), 2.66 (ddd, *J* = 13.4, 10.6, 3.1 Hz, 1H), 2.56 (dd, *J* = 9.0, 6.7 Hz, 2H), 2.27 – 2.12 (m, 7H), 2.04 (dd, *J* = 17.7, 5.3 Hz, 1H), 1.85 – 1.70 (m, 2H), 1.68 (s, 3H), 1.59 (s, 5H), 1.31 (qd, *J* = 7.8, 6.9, 4.3 Hz, 4H), 0.89 (t, *J* = 6.8 Hz, 3H). $^{13}$C NMR (126 MHz, cdcl$_3$) δ 149.61, 147.82, 142.02, 135.75, 132.89, 130.63, 125.88, 124.52, 111.06, 45.61, 38.41, 35.19, 31.47, 30.43, 30.30, 28.68, 23.44, 22.46, 20.95, 19.61, 14.01. IR (FTIR):886, 1032, 1183, 1366, 1770, 2875, 2972 cm-1 MS (ESI): m/z calcd. for $C_{25}H_{34}O_4$ $(M+H)^+$ 399.2529, found 399.2529.

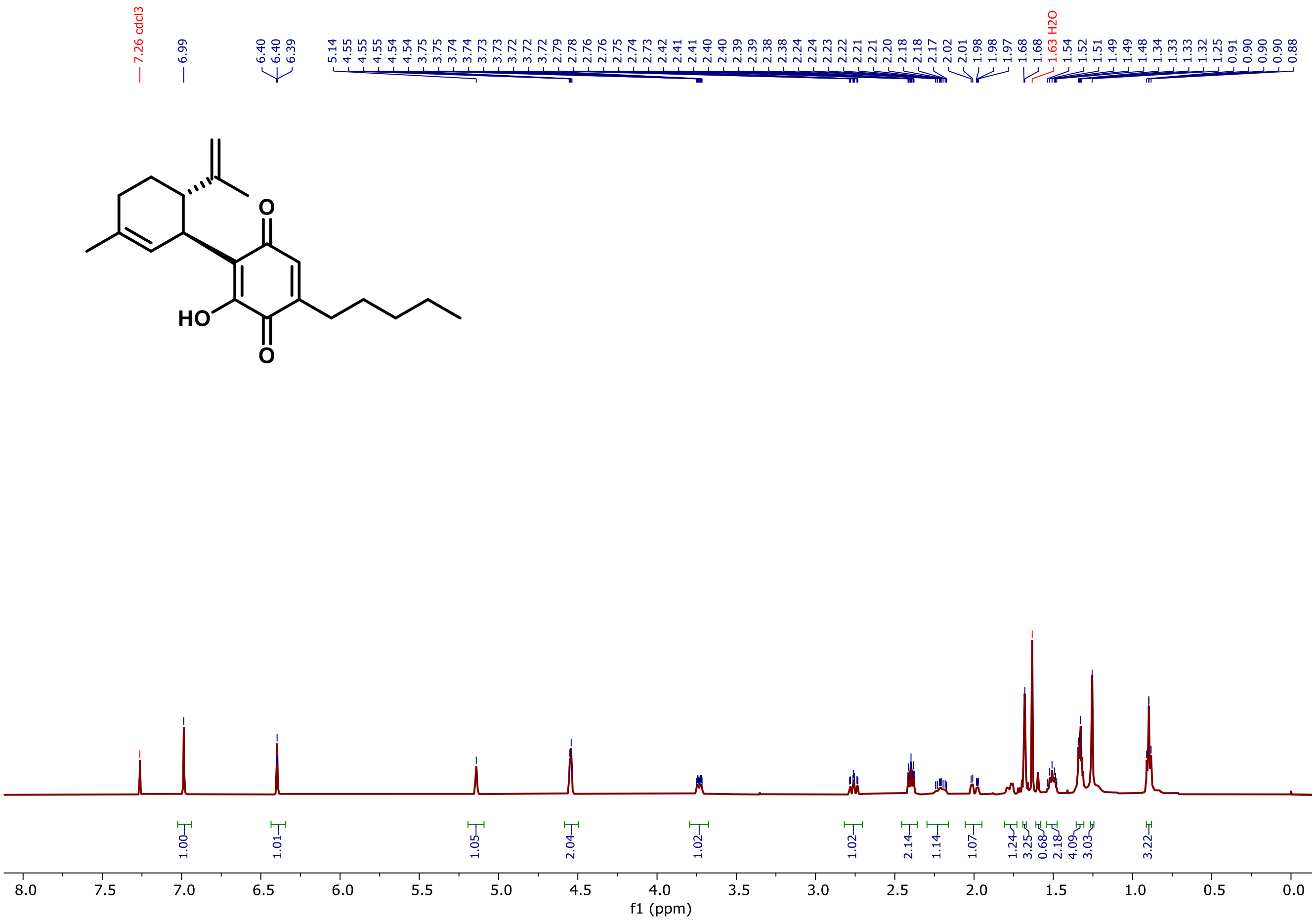

**Figure S1.** $^{1}$H NMR of cannabidiol quinone.

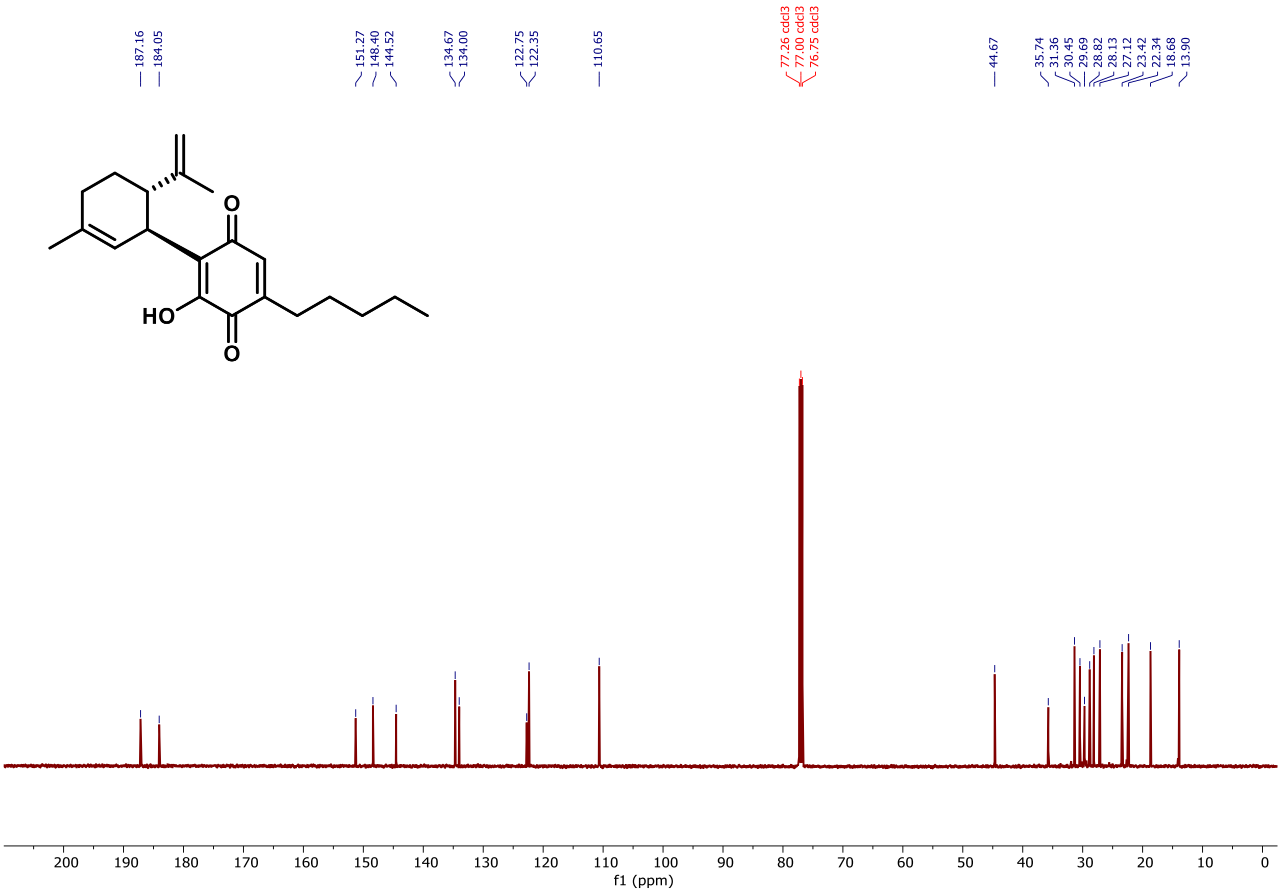


**Figure S2**: $^{13}$C NMR of cannabidiol quinone.

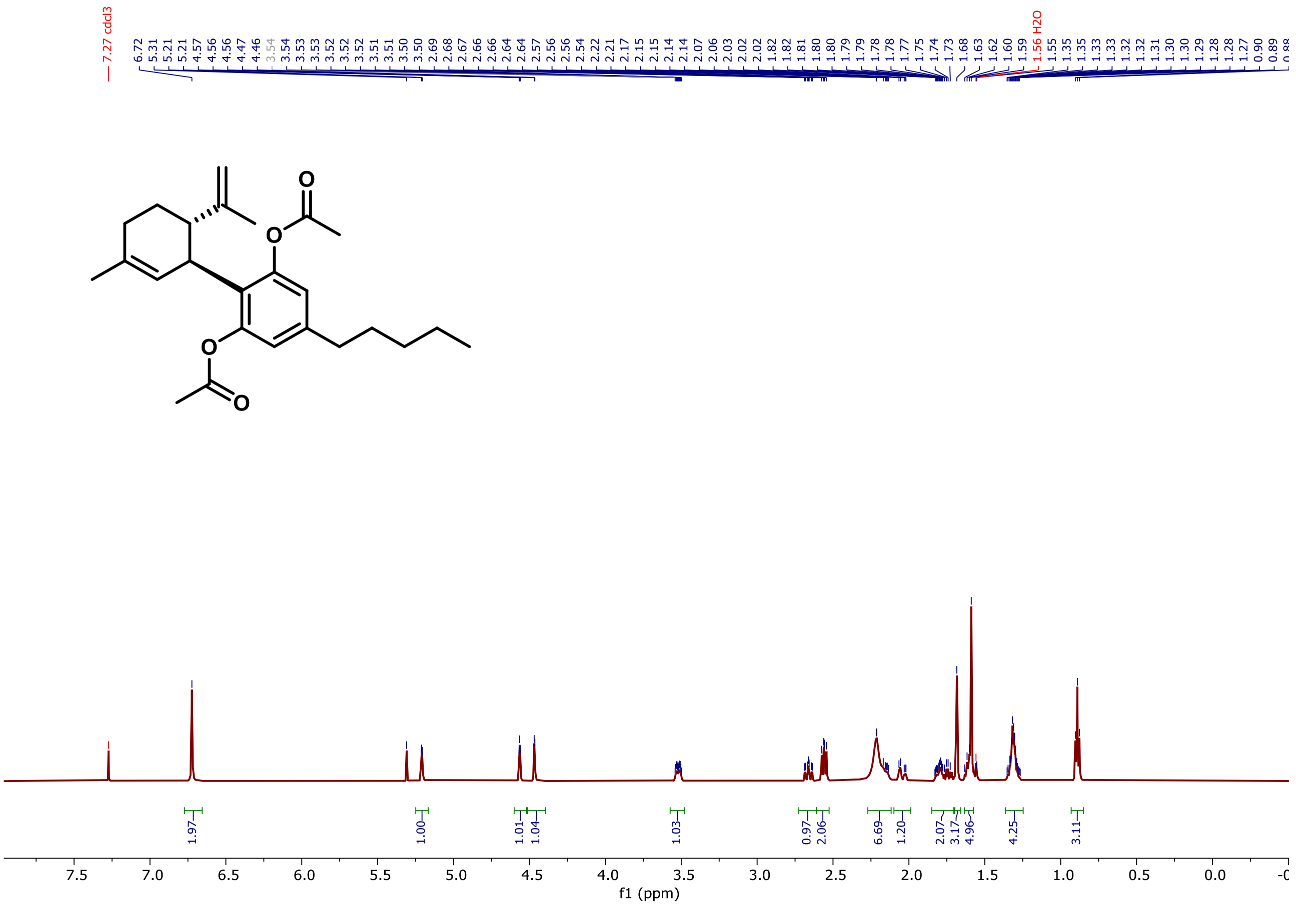

**Figure S3**: $^1$H NMR of diacetyl cannabidiol.

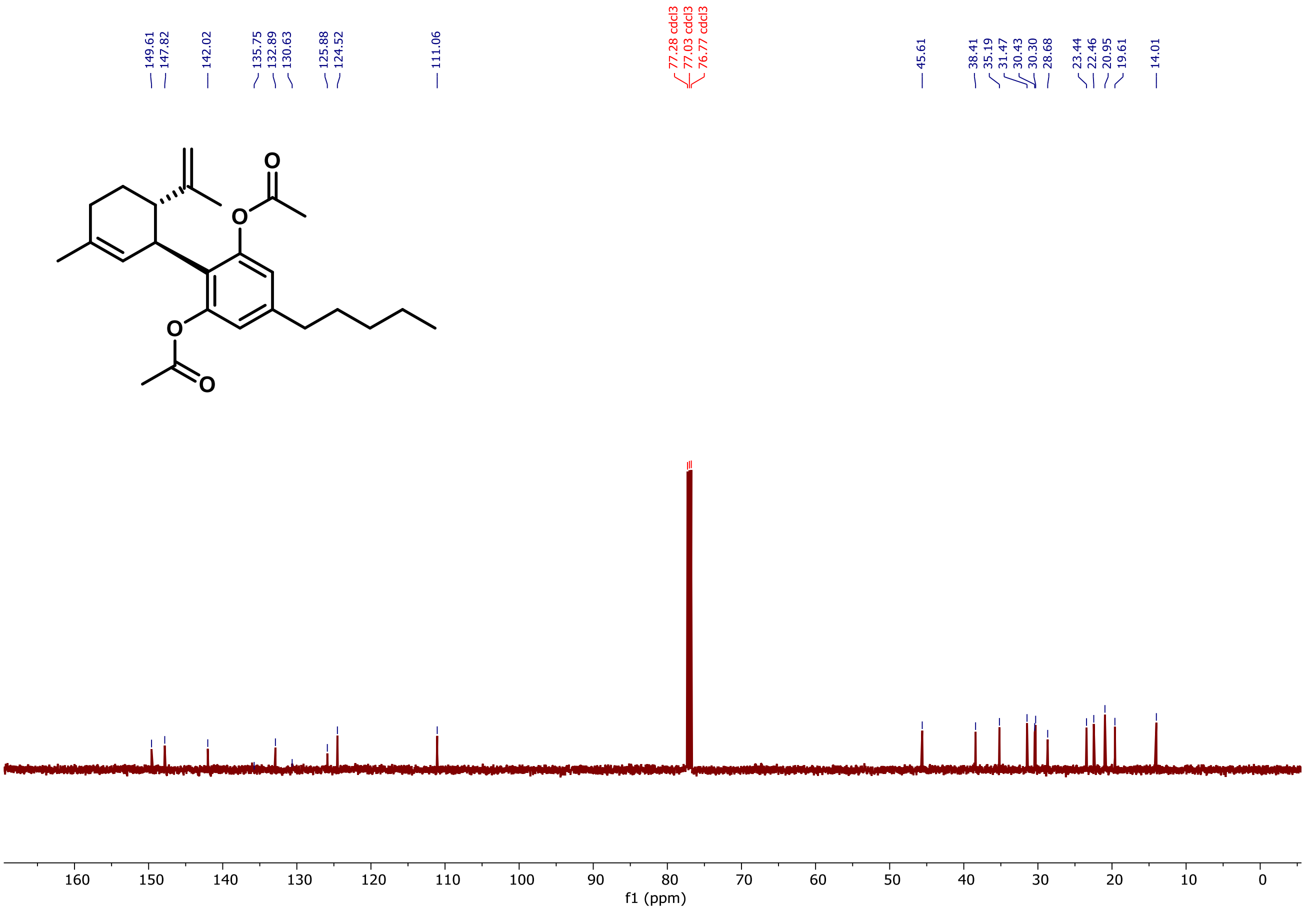


**Figure S4**: $^{13}$C NMR of diacetyl cannabidiol.

### S2. Mass spectrometry

All mass spectrograms were collected with a Thermo Fisher Scientific Q Exactive Plus mass spectrometer with Exactive Tune Software. Ionization was done in ESI Source in positive polarity only. A Hamilton syringe mounted to a syringe pump was used for continuous injection of samples at a 1 mM concentration (CBD in DMSO and CBD in EtOH, both kept for a month) dispensed at a flow rate of 10 µL/min. Each sample ion mass spectra were imported using Xcaliber 4.0 and the relative ion counts of the CBD and CBD-Q normalized and relative abundances calculated.

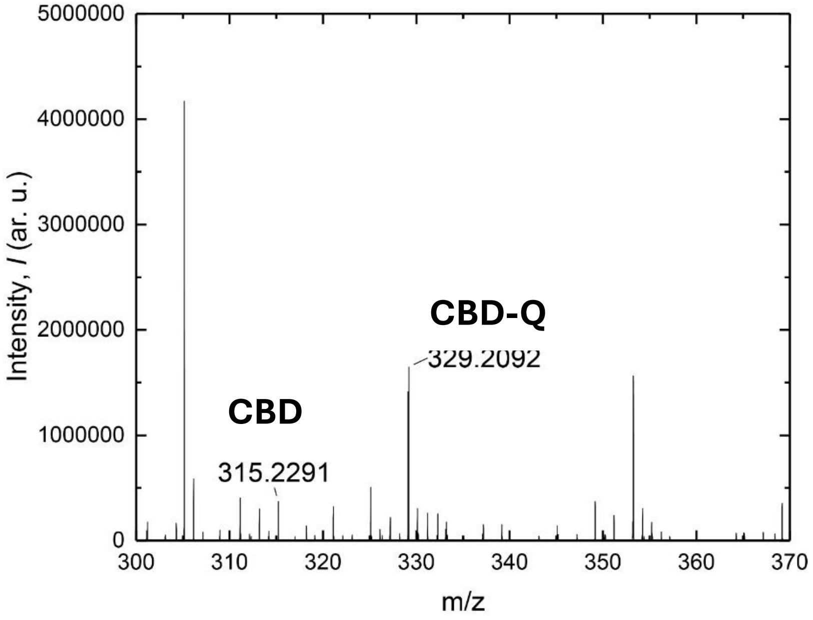


**Figure S5.** Ion spectrum (MS/MS) of CBD in EtOH stored for one month. Observed masses correspond to CBD-Q (*m/z* 329.2092), and CBD (*m/z* 315.2291).

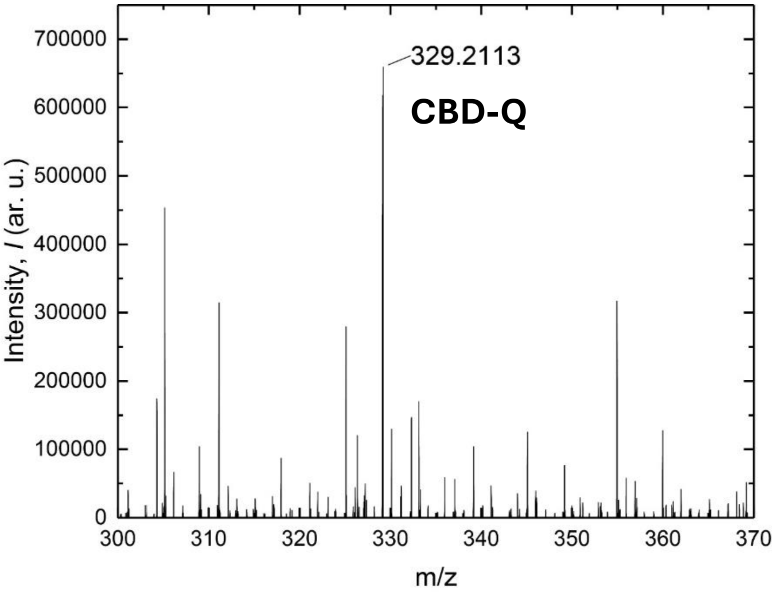


**Figure S6.** Ion Spectrum (MS/MS) of CBD in EtOH stored for one month. Observed mass is CBD-Q (*m/z* 329.2092). No unoxidized CBD was detected.

### S3. Cyclic Voltammetry Procedure

In a borosilicate scintillation cell, tetrabutylammonium hexafluorophosphate supporting electrolyte ($NBu_4PF_6$) solution was added (5 mL, 0.1 M in $CH_3CN$). The glassy carbon working electrode, platinum counter electrode, and platinum pseudo reference electrode were positioned in the cell with ~0.5 cm of the electrode tips submerged in the electrolyte solution. CBD (0.5 mmol) was added to the cell and stirred until the solution was homogenous. A CV spectrum was collected. All spectra were collected with Gamry Instruments Framework, version 6.25, build 3318. All spectral analyses were performed with Gamry Echem Analyst, version 6.25.

#### Cyclic Voltammetry Parameters

$E_{\mathrm{initial}}$ (V): 0.5 vs. $E_{\mathrm{ref}}$

Scan limit 1 (V): 3 vs. $E_{\mathrm{ref}}$

Scan limit 2 (V): 0.4 vs. $E_{\mathrm{ref}}$

$E_{\mathrm{final}}$ (V): 0.4 vs. $E_{\mathrm{ref}}$

Scan rate (mV/s): 200

Electrode area ($cm^2$): 1

Equilibration time (s): 0

*I/E* range mode: auto *I/E* range

Maximum current (mA): 5

Cycles (#): 3

Open circuit (V): 0

Sampling mode: Noise rej

The oxidation potential of CBD dry and purged: 1700 V and 645 mA

The oxidation potential of CBD: 1600 V and 1200 mA

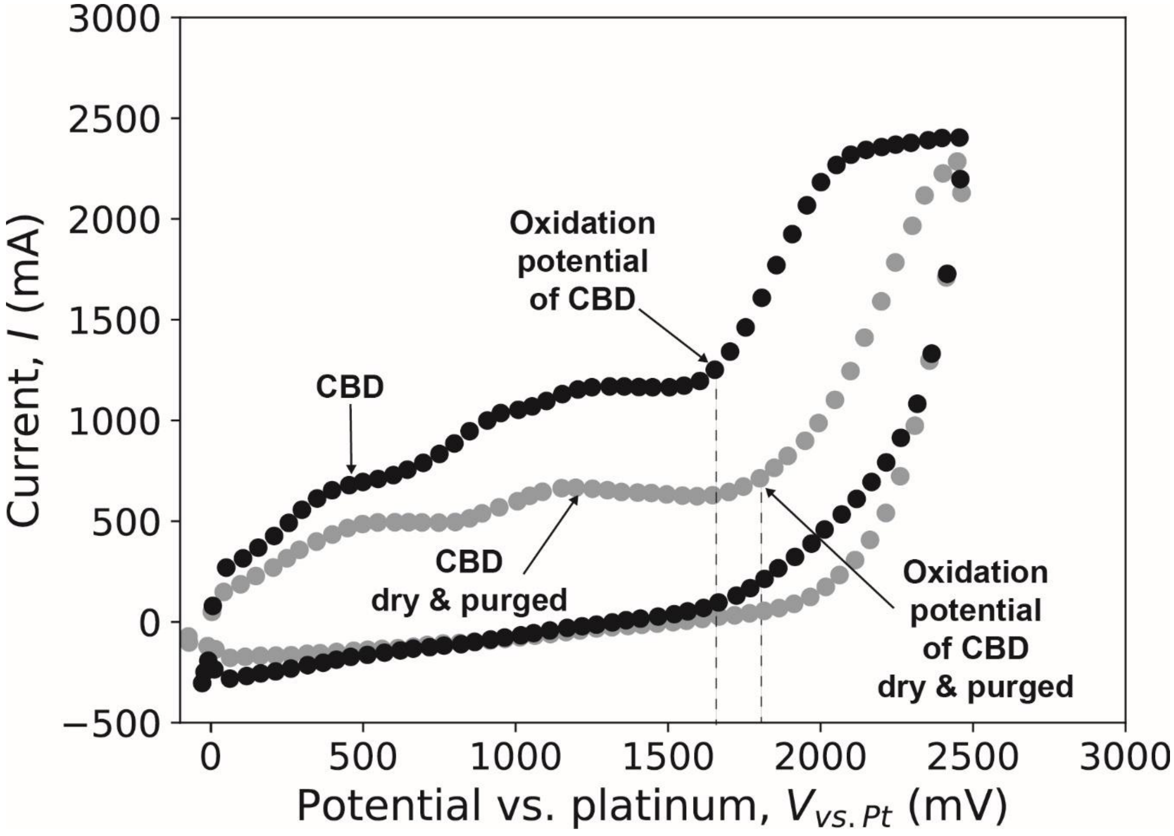


**Figure S7.** Cyclic voltammogram displaying electron flow for CBD in as is electrolyte (black), and dried and purged electrolyte (grey).

**S4. Cell Culture**

***Expanding HUVECs***

Human umbilical vein endothelial cells (HUVECs, Angio-Proteomie cAP-0001) of pass 3, 4, 7, 9 were expanded on 100 mm tissue-culture-treated polystyrene dishes (Corning® 430167) coated with 10 µL/mL fibronectin (FN, Corning® 356008) within a standard cell incubator at 37 ºC, 5% $CO_2$, and 95% humidity until cells were about 70–80% confluent. The HUVECs were fed every other day with the EGM™-2 with BulletKit™ from Lonza (CC-3162), where the media kit already contains about 2% FBS. These four batches of endothelial cells were used for four different external replicate experiments.

***Toxicity Protocol***

The day before the experiment, two 48-well tissue culture-treated polystyrene plates were coated with 10 µL/mL fibronectin (Corning® 354008). To do this, we diluted the stock 1 mg/mL fibronectin with PBS (Gibco™ 70011044) to yield a working concentration of 10 µg/mL. We added 200 µL of the 10 µg/mL fibronectin to each well and then placed in the plate in the cell incubator overnight.

On the day of the experiment, cells were stained with a nuclear stain (Hoechst, ThermoFisher 62249) and a live/dead viability/cytotoxicity kit (Invitrogen L3224), which contained a live cell stain, calcein AM, and a dead cell stain, ethidium homodimer-1. Stock concentrations of stains were as follows: 20 mM Hoechst, 1 mM calcein AM, and 2 mM ethidium homodimer-1. In an aliquot containing 8 mL EGM™-2, 2 µL of each stain was added, then the whole solution vortexed. The final working concentration of each strain was calculated to be 5 µM Hoechst, 0.25 µM calcein AM, and 0.5 µM ethidium homodimer-1. The cells were then rinsed with PBS (Gibco™ 70011044), loaded with the stain-media solution, and placed in the cell incubator for 20 minutes. After 20 minutes, cells were checked for fluorescence and then prepared for passage.

Next, HUVECs were seeded onto the 48-well plates at a seeding density of 10,000 cells per $cm^2$ using standard cell passage protocols. Cells were allowed 3 hrs to settle, and in the meantime, fresh aliquots of the CBD analogs were dissolved to 5 mM stock concentrations in EtOH or DMSO.

We used crystalline CBD sourced from Tree House Hemp, whereas cannabidiol quinone and cannabidiol diacetate were both synthesized in lab with their purities confirmed using mass spectrometry (S1. Synthesis Procedures). Visually, CBD appeared to be a white crystalline powder whereas the CBD quinone and CBD diacetate were both in liquid form with a brown-orange or clear color, respectively. Two vials of each analog were prepped. The crystalline CBD, CBD-Q, and CBD-DA were then dissolved in ethanol (Fisher Scientific BP2818-500) or dimethyl sulfoxide (SIGMA Life Science D2650-100ML) to a stock concentration of 5 mM, then filtered using a 0.1 µm Millex™ PVDF filter (MilliporeSigma™ SLVVR33RS). These 5 mM stock concentrations were diluted further in EGM™-2 to their final working concentrations of 1, 6, or 10 µM to create the loaded media.

After 3 hours of allowing cells to settle, HUVECs were then rinsed with PBS, and each well was fed 300 µL of treated media. Then, the HUVECs were placed in a cell incubator for 24 hours. During the entirety of the experiment, all chemical and biological samples were stored and handled in the dark to the best of our ability (*i.e.*, lights in biohood off, chemical samples covered in aluminum foil).

***Imaging***

After 24 hours, cells were rinsed with PBS and fed a fresh batch of treated media. Next, a widefield epifluorescent inverted microscope (Nikon T2000EU) was used to image cells. Each well was imaged at the center of the well with a 10x magnification objective, 2s exposure, using the microscope's software, Micro-Manager 2.0.0.

The Hoechst stain was excited using a UV light source with 360 ± 40 nm excitation and emission was filtered using a UV Nikon filter (UV-2E/C). The calcein AM stain was excited with blue light with 480 nm ± 30 nm emission and filtered using a blue Nikon filter (B-2E/C). Ethidium homodimer-1 stain was excited using a green light source with 540 nm ± 25 nm emission filtered using a green Nikon filter (G-2E/C).

Each stain was imaged individually as a “slice” at the same focal plane. The three images were compiled together and saved as one image using Micro-Manager 2.0.0. This image was considered the “raw” image.

**S5. Quantitative analysis of cytotoxicity**

For further quantification, we created a counting cells macro to count the cells in each slice of our raw images using the software FIJI ImageJ2. At 10x magnification, for our microscope, 1 pixel was 0.34 μm.

The first part of our counting cells macro converted 16-bit images to 8-bit, subtracted the background, set the threshold, and converted the images to binary. We found that when keeping the images as 16-bit, FIJI had difficulty removing the background for an accurate cell count. Next, a median of radius = 20 pixels was set to eliminate small debris smaller than 7 μm, smaller than the known HUVEC radii of, on average, greater than 20 μm. The analyze particle function had a size exclusion set to 50-infinity pixels (17 μm and larger), around the average known HUVEC diameters. Processed images were visually verified to ensure proper cell count. The resulting *.csv files generated from the analyze particles function were saved automatically and considered our raw *.csv data files. Each raw *.csv file contained all our internal replicates (n = 3) for each experiment date. Eventually, these internal replicates were averaged for each condition in each experiment using Microsoft Excel to generate the external replicates (N = 3 or 4).

For visualization, raw *.csv files from FIJI were then compiled into another data sheet, where calculated averages for the calcein AM stain were compiled. Hoechst stain analysis led to considerable variability, as it stains DNA content of both live and dead cells alike, masking toxicity results. For higher concentrations of CBD analogs, there were higher rates of cell toxicity, resulting in extremely low dead cell attachment, making the ethidium homodimer-1 results unreliable for toxicity quantification. Therefore, the calcein AM stain contained the most information regarding the state of the HUVECs upon exposure to CBD, whereas other slices were variable depending on cell attachment, which could be affected by PBS rinsing.

A Python script was used to graph boxplots, and the significance was added to the plots using Affinity Publisher 2.

All raw data, excel sheets, FIJI macros, and Python scripts are provided as supplemental materials. The following is the pipeline used to generate the graphs, and a guide to the items provided in the supplemental materials:

1. Raw images from fluorescence microscopy

2. Fiji Macro to analyze the raw images: “FijiMacro_countingCells_20240503.ijm”

3. The 4 resulting Fiji Excel Sheet per experimental replicate: “raw” + “YYYYMMDD” + “Summary.csv”

4. Compiled Excel Sheet with all raw data: “compiled_raw.csv”

5. Excel Sheet to specifically average Calcein data: "averagedCalcein_forPython.csv"

6. Excel Sheet with significance calculations: "calcein_Statistics.xlsx"

5. Python script to generate Figure 1, compiled using Jupyter Notebook Version 6.4.12, "averaged_Concentration_Boxplots_20230605" + ".py" or ".ipynb"

6. Used Affinity Publisher 2 to add significance bars.

**S6. Statistical analysis**

After the averages of the internal replicates, n=3, were taken to yield the values of our external replicates, N=3,4, we used Microsoft Excel to conduct an *f*-test between all possible paired combinations of experimental conditions. If the resulting f-test value was less than 0.05, the pair was deemed to have significantly different variances and thus considered a two-sample unequal variance for the *t*-test calculation. If the value were greater than 0.05, then within the *t*-test calculation, the pair would be considered to have a two-sample equal variance.

Afterward, a *t*-test was conducted using Microsoft Excel. Using the averages of the internal replicates, a two-tailed *t*-test was performed. The type (equal or unequal variance) depended on whether the *f*-test yielded equal or unequal variances between the two samples tested. Finally, Microsoft Excel was also used to calculate the standard deviation of each internal replicate, which was then used to calculate the standard error of the mean (SEM).
